# UBC9 and EME1 sumoylation foster ribosomal DNA damage repair in CPT response

**DOI:** 10.1101/2022.04.11.487628

**Authors:** Easa Nagamalleswari, Karishma Bakshi, Jan Breucker, Michaela Gschweitl, Román González-Prieto, Nathalie Eisenhardt, Chronis Fatouros, Viduth K Chaugule, Alfred C.O. Vertegaal, Andrea Pichler

## Abstract

UBC9 sumoylation (S*UBC9) in mammals contributes to SUMO target discrimination, biochemically^1^. Here, we present biological insights by characterizing a sumoylation mimetic UBC9-fusion (^mC^S^~^UBC9) in comparison to its wild-type (^mC^UBC9). We observe that sumoylation promotes UBC9’s stability, nuclear localization and is beneficial for cell survival. We identified EME1, the regulatory subunit of the structure-specific endonuclease EME1-MUS81, as S*UBC9 substrate and demonstrate EME1 and UBC9 sumoylation being advantageous for survival upon Camptothecin (CPT) exposure. Moreover, ^mC^S^~^UBC9 expression enhances double-strand breaks (DSBs) and replication upon CPT treatment, features assigned to the EME1-MUS81 complex^2^. EME1 and ^mC^S^~^UBC9 co-localize in nucleolar repair-condensates, that enlarge and round when exposed to the drug. Together, these findings imply that S*UBC9-induced EME1 sumoylation improves ribosomal rDNA repair, which might prevent dangerous second template switches in highly repetitive ribosomal chromatin and allow the converging replication fork to complete DNA replication. Finally, we discuss implications of these findings for anti-cancer therapy.

## Main

Covalent attachment of the small ubiquitin related modifier (SUMO) to its substrates (sumoylation) involves the hierarchic action of E1, E2 and E3 enzymes. E3 ligating enzymes are supposed to ensure substrate specificity, however 10 bona fide SUMO E3 ligases face more than 6500 identified SUMO substrates and it is unclear how substrate specificity is achieved ^3^. On the basis of biochemical and structural studies, we observed UBC9 sumoylation (S*UBC9) as alternative mechanism to regulate target discrimination, by modifying selected substrates that have a SUMO interaction motif (SIM) in close distance to their modification site^1^. However, the biological significance of mammalian S*UBC9 remained elusive.

Camptothecin (CPT) is a natural topoisomerase I (Top1) inhibitor and its derivatives irinotecan and topotecan are commonly used in clinical anticancer therapy for the treatment of a broad range of solid tumors^4–6^. Top1 resolves supercoiled DNA by introducing single strand breaks (SSB) in one DNA backbone and before religation it remains transiently covalent bound to DNA 3’-phosphate termini, designated as Top1cc. CPT prevents relegation causing Top1ccs and SSBs. These obstacles interfere with transcription and DNA replication and can be converted into single ended double strand breaks (seDSBs) or stalled or reversed replication forks. This causes genomic instability and cell death. However, several cellular DNA repair mechanisms counteract drug treatment, such as, the structure-specific endonuclease complex EME1-MUS81. Upon CPT exposure, EME1-MUS81 generates DSBs, which serve as a prerequisite for replication restart and cell survival^2^. Consistently, both EME1 and MUS81 were among the top ten hits causing resistance to CPT^7^. The balance between damage and repair finally decides the response to therapy. This is reflected in a common adverse feature of CPT-based chemotherapy that varies widely from patient to patient due to acquired or de novo clinical resistance. Ways to overcome this resistance would greatly improve therapy and allow the definition of personalized treatments. The molecular mechanisms leading to resistance are poorly understood.

Connections between the SUMO system and cancer progression are well established^8–14^. Cancer drug resistance, tumor metastasis and relapse correlate with constitutive high levels of individual SUMO enzymes or global sumoylation levels^15,16^. Inhibition of the SUMO E1 enzyme delays tumor progression by decreasing cancer cell viability in cellular and xenograft model systems^9,10,12,17^ and overexpression of a dominant negative UBC9 mutant has been shown to improve response to the CPT derivative topotecan^18^.

In the presented study, we investigated the biological function of UBC9 sumoylation upon CPT treatment and identified EME1 as substrate. Moreover, we found that sumoylation of both, UBC9 and EME1, promoted cell survival upon CPT exposure by inducing DSBs and replication in the nucleolus. Finally, we discuss a model how S*UBC9 induced EME1 sumoylation prevents genome instability in the highly repetitive ribosomal chromatin, disclosing molecular insights in the regulation of response to cancer treatments.

### A SUMO fusion can mimic

Sumoylation is reversible and usually very transient due to cellular SUMO proteases. This, together with the fact that mutation of major sumoylation sites often lead to modification of minor sites, makes it challenging to study the consequences of substrate sumoylation. *In vitro,* mammalian UBC9 is mainly modified at Lys14^1^. Mutation of this lysine (UBC9-K14R) resulted in modification of Lys153. When we also mutated this lysine (UBC9-K14/153R) we detected modification at Lys18, Lys49, Lys146 (^1^ and unpublished observations). Detection of UBC9 sumoylation at Lys14 in cells is difficult as conventional mass spectrometric approaches, to map SUMO conjugation sites, use trypsin digestion that cleaves the Lys14-containing UBC9 peptide so small that it cannot be specifically assigned. Moreover, UBC9 interacts with SUMO in various manners: (1) for its catalytic activity, it forms a thioester bond via Cys93, (2) it non-covalently binds SUMO via its backside, opposite to the catalytic cleft and (3) it is sumoylated via an isopeptide linkage mainly on Lys14^3^. To overcome all these technical hurdles and to distinguish between various UBC9 functions, we decided to investigate, whether a gain-of-function mutant, by fusing a non-cleavable SUMO N-terminal to UBC9 (S^~^UBC9, peptide bond linked SUMO), can mimic SUMO conjugation of UBC9 (S*UBC9, isopeptide bond linked SUMO). Specifically, we compared bacterial expressed, *in vitro* sumo(2)ylated UBC9 (S*UBC9)^1^ and a linear SUMO2^~^UBC9 fusion (S^~^UBC9) with unmodified UBC9 for their abilities to modify the established substrate SP100. As shown in Fig. 1a, both UBC9 sumoylation variants follow the same trend and significantly enhance modification of SP100 compared to unmodified UBC9.

**Fig. 1.**
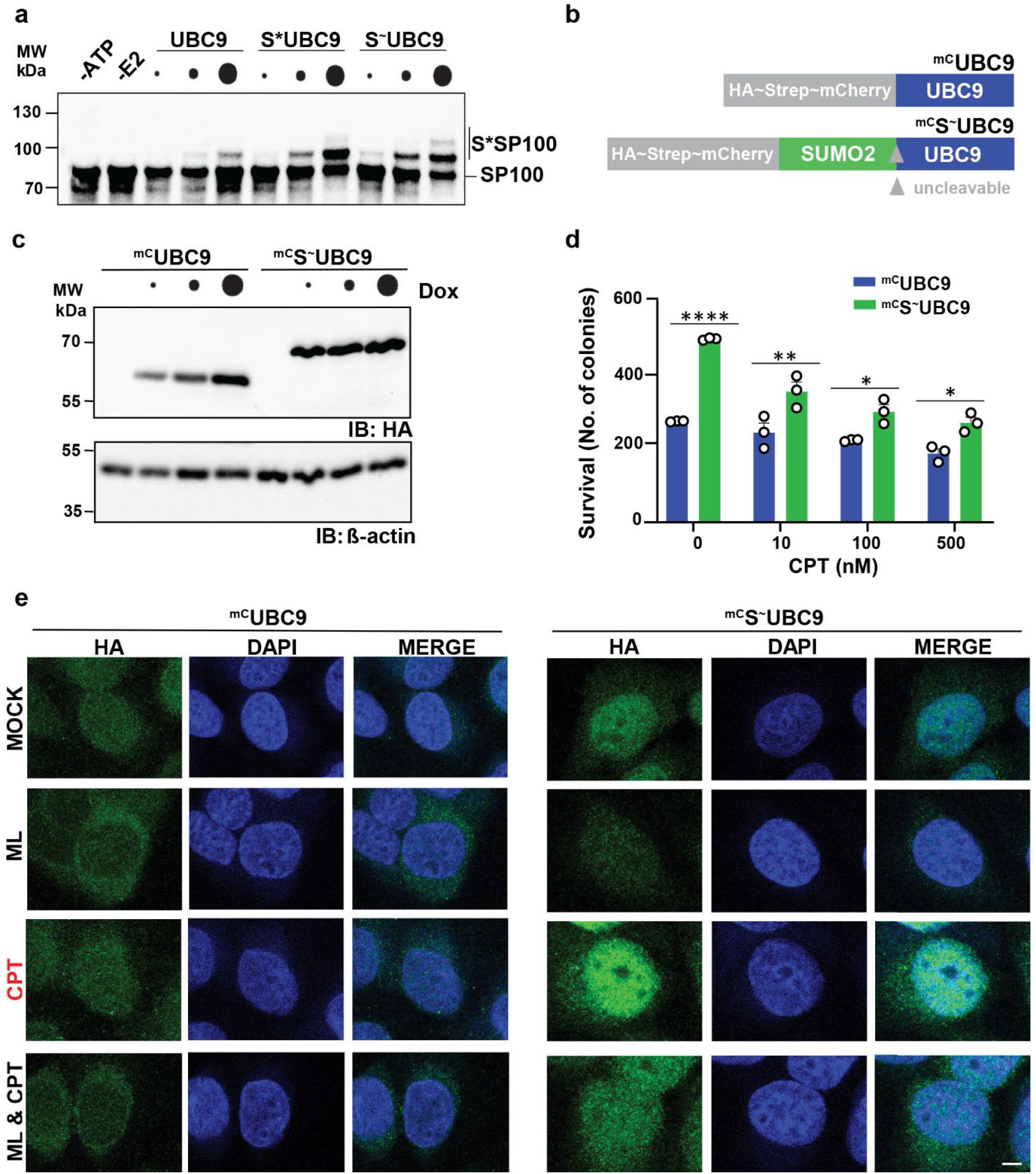
Sumoylation promotes UBC9’s stability, nuclear localization and cell survival. **(a)** Immunoblot of a multi-turnover *in vitro* sumoylation assay. Increasing concentrations (0, 10, 50 or 250 nM) of UBC9, S^~^UBC9 or S*UBC9 were incubated with 200 nM GST-SP100, 60 nM E1, 2 μM SUMO2 amd 5 mM ATP in 20 μl reactions 30 minutes at 30 °C. GST-SP100 sumoylation was monitored after 30 minutes at 30 °C. **(b)** Schematic representation of HA-Strep-mCherry tagged full-length UBC9-variants: ^mC^UBC9 wild-type and the SUMO2 fusion ^mC^S^~^UBC9, that were stable integrated in the FRT locus of U2OS cells for doxycycline (Dox) inducible expression. **(c)** Immunoblot showing expression levels of stable integrated ^mC^UBC9 and ^mC^S^~^UBC9 induced by different (Dox) concentrations (0, 10, 40, 160 ng/ml) for 16 hours. Cell lysates were resolved on SDS -PAGE and UBC9-variants expression (anti-HA) was monitored in comparison to β-actin levels. **(d)** Diagram of clonogenic survival assay. ^mC^UBC9 or ^mC^S^~^UBC9 expression was induced for 16 hours with 30 and 10 ng/ml Dox, respectively, before cells were treated with indicated concentrations (0, 10, 100 and 500 nM) of CPT for one hour. Colonies were stained with 0.5% Crystal Violet after 7-12 days and analyzed with ImageJ. Bars show the average (+SEM) surviving colonies of three different experiments performed in triplicate and are significant based on two-way ANOVA Tukey test by GraphPad Prism analysis (*: p-value < 0.05; **: p-value < 0.01, ****: p-value< 0.0001). Graphs with plotted values normalized to mock conditions are shown in Extended Data Fig. 1a. **(e)** Immunofluorescence analysis of ^mC^UBC9 and ^mC^S^~^UBC9 upon different treatments. Expression of ^mC^UBC9-variants as in (D) before treatment with either DMSO (vehicle control, MOCK), ML (ML792, 1 μM for 2 hours), CPT (10 μM for 1 hour) or ML & CPT. Co-staining was with anti-HA antibodies (labelled with Alexa 488, green) for detecting UBC9-variants and DAPI (blue) for visualization of nuclei. Shown are individual stains and merges of representative single cells, according multi cell images are shown in Extended Data Fig.1b. Scale bar represents 10 μm.

From these data we conclude that a N-terminal linear S^~^UBC9 fusion can be studied as a proxy of a S*UBC9 conjugate.

### ^mC^S^~^UBC9 is stabilized and promotes cell survival

To understand consequences of UBC9 sumoylation, we generated doxycycline (dox) inducible U2OS cell lines expressing HA-Strep-mCherry tagged (^mC^) UBC9-variants: ^mC^UBC9, as wild-type and the SUMO2 fusion ^mC^S^~^UBC9 (Fig. 1b). Titration of Dox induced expression indicates that ^mC^S^~^UBC9 is stabilized compared to ^mC^UBC9 (Fig. 1c). Sumoylation has key functions in DNA damage repair and overexpression of a dominant negative UBC9 mutant increased sensitivity to the CPT derivative topotecan^18^. Therefore, we compared the two UBC9 variant-expressing cell lines in terms of clonogenic survival upon treatment with different concentrations of CPT. Fig. 1d shows the average percentage of surviving colonies of three biological replicates performed in triplicates. Values normalized to untreated conditions are shown in Extended Data Fig. 1a. Already under untreated conditions, but also at lower CPT concentrations tested (10 and 100 nM), ^mC^S2~UBC9 expression is beneficial to cell survival and colony formation.

### S^~^UBC9 is enriched in the nucleus

We then examined the intracellular localization of the two UBC9-variants by confocal microscopy (Fig. 1e and Extended Data Fig. 1b). Throughout the manuscript representative single-cell images are shown in Main Figures, corresponding multi-cell images in Extended Data Figures. In mock-treated cells, both variants were localized throughout the cell, with ^mC^S^~^UBC9 showing enrichment in the nucleus (first panel). When we exposed the cells to the SUMO-E1 inhibitor ML972 (ML) for two hours before analysis, ^mC^UBC9 was mostly in the cytosol, whereas ^mC^S^~^UBC9 remained in the nucleus but was less abundant (second panel).

These data indicate that sumoylation enriches UBC9 in the nucleus. Next, we treated the cells with CPT for one hour and observed that both forms accumulated in the nucleus, with ^mC^S2~UBC9 showing the brightest staining (third panel). Pretreatment with ML792 one hour before the addition of CPT abolished nuclear localization of ^mC^UBC9 and resulted in a reduced nuclear signal from ^mC^S2~UBC9 (fourth panel). These data suggest that UBC9 enters the nucleus as activated SUMO thioester charged or sumoylated form. Stress treatment increases nuclear localization depending on its activity.

In summary, sumoylation enhances UBC9’s stability, nuclear localization and provides a survival benefit in mock and CPT exposed cells.

### EME1 is a substrate for S*UBC9

To identify putative substrates of S*UBC9 that might mediate the survival benefit, we relied on a screen in which we performed *in vitro* sumoylation reactions on a human protein array with either unmodified (UBC9) or sumo(1)ylated UBC9 (S1*UBC9) as outlined in Fig. 2a. We identified substrates whose sumoylation state remained enhanced, reduced, or unchanged in the presence of S1*UBC9 compared to UBC9. Among the enhanced sumoylated candidates, we searched for proteins with demonstrated functions in CPT repair and found EME1, the regulatory subunit of the structure-specific endonuclease EME1-MUS81 complex^2,7^. We continued by verifying EME1 as an actually enhanced substrate of both S2*UBC9 and S2^~^UBC9 in *in vitro* sumoylation assays (Fig. 2b). However, when we turned to E3 ligases, we found that several SUMO E3 ligases also promoted EME1 modification *in vitro,* but to varying degrees (Fig. 2c). Of note, we found no evidence that S*UBC9 enhanced the activity of E3 ligase in *in vitro* sumoylation assays (Fig. S2A and^19^). In our previous work, we proposed that substrates for efficient modification with S*UBC9 have a SUMO consensus (SCM: ΨKxE, where Ψ is a large hydrophobic amino acid and X is any amino acid) and a SUMO interaction motif (SIM) in close distance, as outlined in the cartoon in Fig. 2d^1^. Actually, we found one SCM and two putative SIMs in the N-terminus of EME1 (Fig. 2e). Mutation of the key residues of these motifs confirmed that EME1 lysine 27 is the major sumoylation site and that SIM1, but not SIM2, is required for modification with S2*UBC9 (Fig. 2f).

**Fig. 2.**
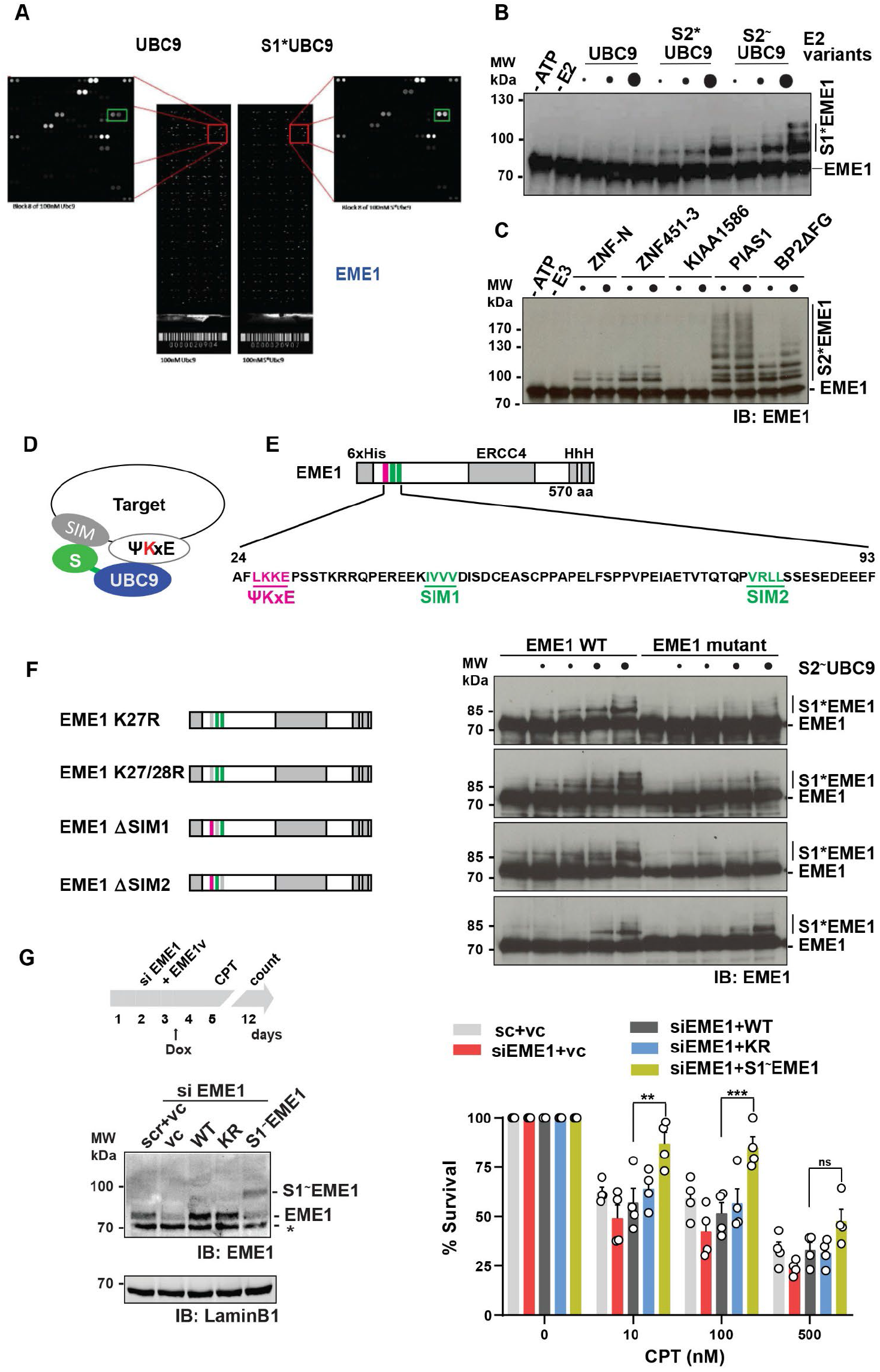
EME1 is a S*UBC9 substrate and its sumoylation also promotes cell survival. **(a)** Shown are slides of the in vitro sumoylation reaction on the protein array that led to the identification of EME1 as an enhanced substrate for S*UBC9. Rectangle shows enhanced substrate sumoylation in the presence of S*UBC9 in comparison to UBC9 as we observed for EME1. *In vitro* sumoylation reactions were performed on protein array slides with 4 μM HA-SUMO1, 100 nM E1, an ATP regenerating system and either UBC9 or S1*UBC9 at two concentrations each, 100 nM or 500 nM. Incubation was for one hour at 30°C. Detection was with anti-HA antibody and Alexa647 labelled secondary antibodies. **(b)** Immunoblot of E2 dependent *in vitro* sumoylation of EME1 with increasing concentrations of UBC9, S2*UBC9 or S2^~^UBC9 (125, 250 and 500 nM) in the presence of 90 nM Aos1-Uba2, 200 nM MUS81-EME1 and 4.4 μM SUMO1. EME1 sumoylation was monitored after 60 minutes at 30 °C. **(c)** Immunoblot of *in vitro* sumoylation assays with SUMO2 to measure E3 ligases dependent sumoylation of EME1. Analyzed were two different E3 ligase concentrations of ZNF-N, ZNF451-3 and KIAA1568 at 60 and 180 nM, PIAS1 at 40 and 160 nM and BP2ΔFG at 4 and 20 nM, in presence of 50 nM Aos1-Uba2, 25 nM UBC9, 200 nM EME1 and 2 μM SUMO. EME1 sumoylation was monitored after 60 minutes at 30 °C. **(d)** Schematic representation of the interaction between S*UBC9 and a SUMO consensus site and SIM containing target protein.**(e)** The EME1 N-terminus bears a SUMO consensus site around Lys27 (red) and two putative SIM motifs (green). Schematic representation and the amino acid sequence of the N-terminus is shown. **(f)** Cartoon presentation of EME1 mutations (left panel). Mutations are shown in gray. Immunoblot of E2 dependent *in vitro* sumoylation of indicated EME1 mutants compared to EME1-WT with increasing concentrations of S2^~^UBC9 (0, 62.5, 125, 250 and 500 nM) in the presence of 90 nM Aos1-Uba2, 200 nM MUS81-EME1 and 4.4 μM SUMO1(right panel). EME1 sumoylation was monitored after 60 minutes at 30 °C. **(g)** Schematic outline of the clonogenic survival assay (upper panel). HeLa cells were seeded on day 1, siRNA EME1 or a scrambled siRNA (sc) knock down was performed on day 2. On day 3 the cells were transfected with doxycycline (Dox) inducible expression plasmids for EME1-WT (WT), EME1-K27/28R (KR), S1^~^EME1 linear fusion or a vector control (vc) and doxycycline (final concentration 1 ng/ml) was added in the evening of the same day. On day 5 the cells were treated for 1 hour with different CPT concentrations (0.01, 0.1, 0.5 and 1 μM) or DMSO. Subsequently, the cells were allowed to recover for one week. Surviving colonies were fixed, stained and counted. Immunoblots with indicated antibodies are shown from day 5 before CPT treatment (Left lower panel). Bar charts depict the average values and error bars (+SEM) of four different experiments performed in triplicate (Right panel). Values were normalized using untreated conditions colony counts as 100%. Differences between wild-type and S1^~^EME1 constructs are significant based on two-way ANOVA followed by Tukey’s test for multiple comparisons (**: p-value<0.01; ***: p-value<0.005; ns: not significant).

With these data, EME1 was demonstrated to be a novel substrate for S2*UBC9 *in vitro.*

### S^~^EME1 also promotes cell survival

We next examined whether sumoylation of EME1 exhibits a similar phenotype to S*UBC9 upon CPT treatment. To address this question, we performed clonogenic survival assays in HeLa cells after RNAi mediated depletion of endogenous EME1 and replacement with EME1-variants (EME1-v): EME1-WT (wild-type), a loss of function EME1-KR (EME1 K27/28) mutant, a sumoylation mimetic S1^~^EME1 N-terminal fusion, and the empty vector served as replacements. MUS81 was co-expressed to ensure expression of a functional complex. We used a Dox inducible system to achieve expression levels comparable to those of endogenous EME1. However, for S1^~^EME1 we were unable to obtain higher expression levels in transient expressions. Fig. 2g shows a cartoon of the experimental procedure (upper left panel), a representative example of the replacements (lower left panel) and the average percentage of surviving colonies from five biological replicates performed in triplicate (right panel). As described for MUS81^2^, we observed that depletion of EME1 also resulted in reduced survival at all CPT concentrations tested. Survival was brought to almost similar levels with EME1-WT and also with EME1-KR. Though, with S1^~^EME1, we observed a significantly increased survival rate, comparable to the phenotype observed with S2^~^UBC9. Subsequent analyses showed that EME1-KR is still efficiently modified in cells (Extended Data Fig. 2b) and that modification jumps to minor sites when the major site is mutated in E3-dependent *in vitro* sumoylation reactions (Extended Data Fig. 2c). These observations strengthen the notion that an artificial N-terminal SUMO fusion can mimic natural sumoylation and the exact position of EME1 sumoylation appears irrelevant.

In summary, we demonstrated EME1 as substrate for S*UBC9 and showed that sumoylation of both proteins is beneficial to cell survival.

### S^~^UBC9 expression induces DSBs and replication, properties that have been demonstrated for EME1-MUS81

The EME1-MUS81 complex is involved in CPT-induced DNA damage repair by inducing DSBs as a prerequisite for replication restart^2^. If EME1 sumoylation is regulated by UBC9 sumoylation in cells, expression of ^mC^S^~^UBC9 should also lead to an increase in DSBs and replication. We addressed this question by examining ^mC^UBC9-variant expressing cells for their ability to induce γH2AX (Serine139-phosphorylated histone H2AX) as marker for DSBs (Fig. 3a) and to incorporate EdU (5-ethynyl-2’-deoxyuridine) as indicator of replication efficiency (Fig. 3b) and measured the corresponding fluorescence signals by Fluorescence-Activated Cell Sorting (FACS) analysis. We compared mock, CPT and ML792 & CPT treated cells and found that expression of ^mC^S^~^UBC9 increased both DSB formation (Fig. 3a) and replication (Fig. 3b) under all conditions. However, this was only partially reversed by pretreatment with ML792, suggesting sumoylation-dependent and -independent functions of ^mC^S^~^UBC9.

**Fig. 3.**
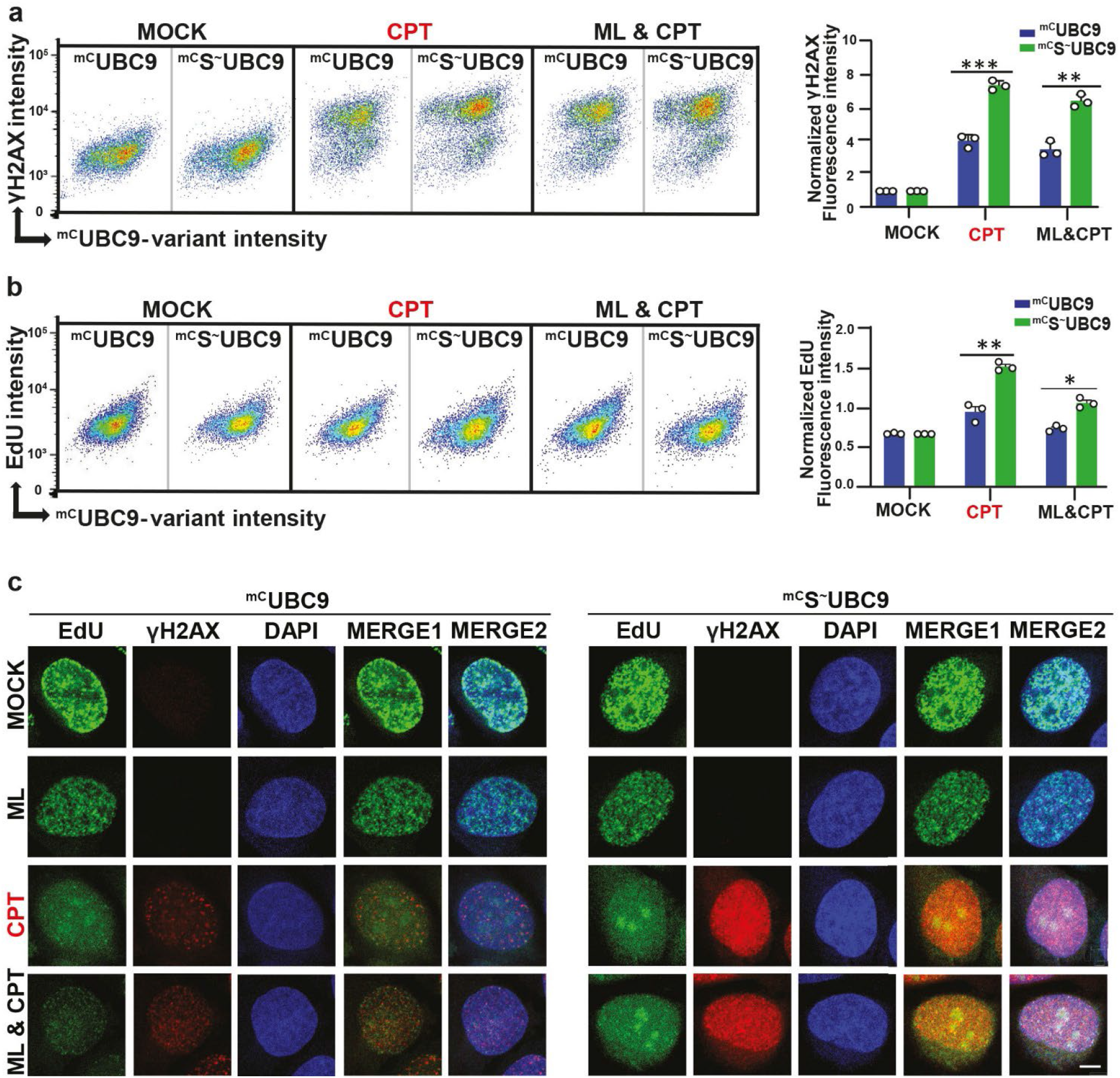
S^~^UBC9 expression induces DSBs and replication upon CPT exposure. **(a)** Quantitative analysis of DNA damage upon CPT exposure. Asynchronously grown population of U2OS cells induced for indicated ^mC^UBC9-variants expression before treatment with DMSO (MOCK), CPT (10 μM for 1 hour) or ML & CPT (ML792, 1 μM for 2 hours) in the presence of 100 μM EdU for 1 hour, fixed and stained with DAPI, γH2AX, and EdU. Scatter plots (left panel) depict cell populations with each dot presenting a single cell. γH2AX with error bars (+SEM) of γH2AX fluorescent intensities (anti-γH2AX coupled Alexa 647) are shown as function of UBC9-variant fluorescent intensities (anti-mCherry coupled to Alexa 594). Bar charts (right panel) show average values and error bars (+SEM) of γH2AX levels of three different experiments performed in triplicate for the indicated cell line and condition. Values were normalized to mock conditions. Differences between ^mC^UBC9 or ^mC^S^~^UBC9 expressing cells are significant based on two-way ANOVA Tukey test by GraphPad Prism analysis (*: p-value < 0.05; **: p-value < 0.01, ***: p-value < 0.001). **(b)** as (a) but quantitative analysis of replication upon CPT exposure. Scatter blots depict EdU signals (coupled to Alexa488) as function of UBC9-variant fluorescent intensities (anti-mCherry coupled to Alexa 594) and bar charts show average values and error bars (+SEM) of EdU levels from three different experiments in triplicates performed for the indicated cell line and condition. **(c)** Immunofluorescence analysis to monitor DSBs (γH2AX) and replication (EdU) in ^mC^UBC9 and ^mC^S^~^UBC9 expressing cells upon indicated treatments. ^mC^UBC9 or ^mC^S^~^UBC9 expression was induced before treatment with DMSO (MOCK), ML (ML792, 1 μM for 2 hours), CPT (10 μM for 1 hour) or ML & CPT in the presence of 100 μM EdU for 1 hour. Confocal imaging was performed after detecting EdU incorporation with Click-iT™ Plus EdU Kit (coupled to Alexa 488, green) and staining with anti-γH2AX antibodies (coupled to Alexa 594, red), respectively. DAPI (blue) was used for visualization of nuclei. Green and red fluorescence shows localization of EdU (replication) and γH2AX (DSB) respectively, merge 1 shows co-localization of EdU and γH2AX, and merge 2 shows co-localization of proteins along with DAPI. Shown are representative single cells, according multi cell images in Extended Data Fig. 3a. Scale bar represents 10 μm.

Next, we compared DNA replication (EdU) and DSBs (γH2AX) in both cell lines by confocal microscopy using the same treatments. As shown in Fig. 3c and Extended Data Fig. 3, EdU staining in both cell lines shifts from a rather small punctate staining in cells mock and ML792-treated to a more diffuse staining with enrichment in large nucleoli resembling speckles in cells upon CPT treatment. These condensates are brighter and respond to ML792 pretreatment in ^mC^UBC9 but not in ^mC^S^~^UBC9 expressing cells, suggesting a role for UBC9 sumoylation in this particular cellular compartment. γH2AX staining was strongly induced by CPT in both cell lines but brighter in ^mC^S^~^UBC9 lines.

Taken together, upon CPT exposure ^mC^S^~^UBC9 enhances DSBs and replication, features attributed to EME1-MUS81. Moreover, replication in ^mC^S^~^UBC9 expressing cells appears to be strongly enriched in nucleoli in a CPT-dependent manner.

### S^~^UBC9 is enriched in the nucleolus

Cellular DSBs lead to the recruitment of various DNA repair factors to insoluble structures that can be visualized by treating cells with a detergent and sucrose mixture to release soluble proteins before immunofluorescence staining^20,21^. We applied this technique to gain insight into whether we can enrich UBC9-variants in such repair structures and whether we can detect changes upon treatment with CPT and ML792. As shown in Fig. 4a and Extended Data Fig. 4a, this was indeed the case. CSK buffer treatments enriched both ^mC^UBC9-variants at the nuclear envelope and in nuclear repair condensates that again resembled nucleoli. The recruitment of UBC9 to the nuclear envelope is well established by its interaction with the SUMO-E3 ligase RanBP2^22^ and appeared comparable between ^mC^UBC9-variants. The nature of the nuclear condensates differed between UBC9-variants in number, size and shape. While speckles appeared small and diffuse in mock treated ^mC^UBC9 expressing cells, ^mC^S^~^UBC9 expression caused enrichment in a few larger nucleoli resembling condensates. ML792 treatment significantly reduced speckle formation, suggesting the involvement of additional sumoylation events. CPT exposure increased the intensity and the shape of these condensates, especially in ^mC^S^~^UBC9 expressing cells. CPT induced condensates were less sensitive to ML792 treatment in ^mC^S^~^UBC9 than in ^mC^UBC9 expressing cells. Again, these condensates resembled nucleoli. Because nucleoli have coordinating functions in stress response, such as DNA damage, we investigated whether these larger speckles actually represent nucleoli by performing co-localization experiments with the nucleolar marker SYTO RNA Select Green Fluorescent Cell Stain in ^mC^S^~^UBC9 expressing cells. Our data confirmed the co-localization in both mock and CPT treated cells (Fig. 4b and Extended Data Fig. 4b). Again, CPT treatment resulted in increased accumulation in the nucleoli, accompanied by a change in the shape of the condensates from fuzzy to round.

**Fig. 4.**
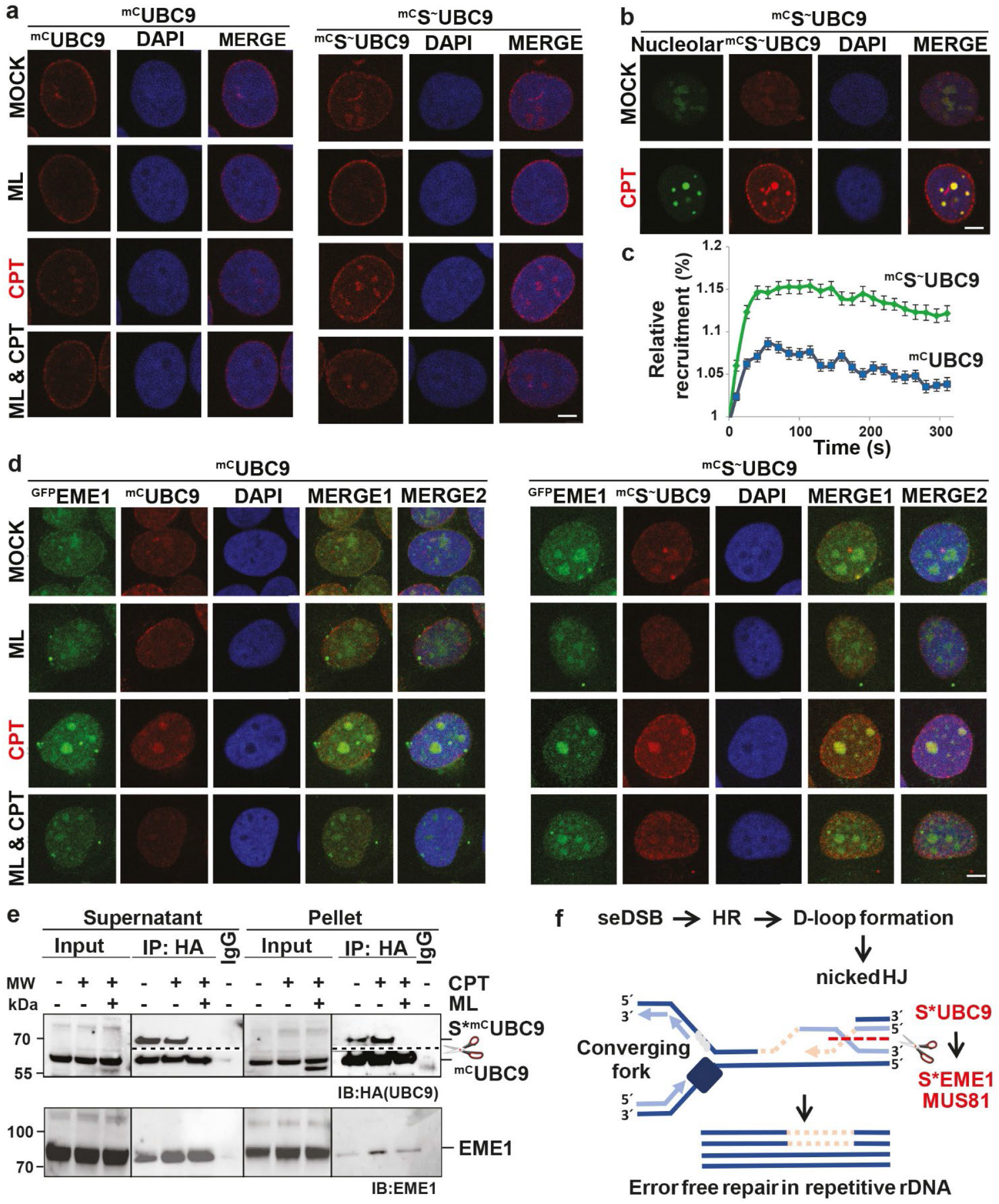
S^~^UBC9 and EME1 co-localize in the nucleolus. **(a)** Immunofluorescence analysis to monitor ^mC^UBC9-variant localization in CSK-buffer insoluble cellular condensates. ^mC^UBC9 or ^mC^S^~^UBC9 expression was induced before treatment with DMSO (MOCK), ML (ML792, 1 μM for 2 hours), CPT (10 μM for 1 hour) or ML & CPT, followed by CSK-buffer treatment to release soluble cellular proteins. After fixation, cells were stained with anti-mCherry antibodies coupled to Alexa 594. DAPI (blue) was used for visualization of nuclei. mCherry signals (red) show intra-nuclear localization of ^mC^UBC9 and ^mC^S^~^UBC9, and merge shows localization of proteins along with DAPI. Shown are representative single cells, according multi cell images in Extended Data Fig. 4c. Scale bar represents 10 μm.**(b)** Immunofluorescence analysis to investigate nucleolar localization of ^mC^S^~^UBC9. ^mC^S^~^UBC9 expression was induced, treated with DMSO (MOCK) or CPT before CSK buffer incubation and fixation. Staining was with 5μM Syto RNA select green fluorescent nucleolar stain (green), DAPI (blue) for visualization of nuclei and mCherry (red) for ^mC^S^~^UBC9 detection, nucleolar stain. Shown are individual stains and merges (^mC^S^~^UBC9 with nucleolar stain) of representative single cells, according multi cell images in Extended Data Fig.4b. Scale bar represents 10 μm. **(c)** Recruitment of ^mC^UBC9-variants to DNA damage tracks. ^mC^UBC9 and ^mC^S^~^UBC9 expression was induced, micro-irradiated with a 2-photon laser and recruitment of the tagged protein was followed in a time course. Shown is the relative recruitment quantification of the ^mC^UBC9 constructs to the laser induced DNA damage tracks. Average and SEMs of three different experiments are shown (N ^mC^UBC9=48; N ^mC^S^~^UBC9=62). **(d)** Co-localization of ^GFP^EME1 with ^mC^UBC9-variants. As in (a) but cell lines were in addition transfected with ^GFP^EME1that was visualized by anti-GFP coupled to Alexa 488 (green). Shown are individual stains (^GFP^EME1 in green, ^mC^UBC9-variants in red, DAPI in blue) and merges (MERGE1: ^GFP^EME1 with ^mC^UBC9-variants, MERGE2: ^GFP^EME1 with ^mC^UBC9-variants and DAPI) of representative single cells, according multi cell images in Extended Data Fig.4c. Scale bar represents 10 μm. **(e)** ^mC^UBC9 sumoylation and EME1 interaction monitored by immunoprecipitation (IP). Induction of ^mC^UBC9 expression and drug treatments as in (a). Cells were lysed in RIPA buffer, sonicated and separated in supernatant and pellet fractions. Pellets were dissolved in RIPA buffer containing 0.5 % SDS and after lysis diluted to 0.1 % SDS. ^mC^UBC9 IPs were performed with anti-HA antibodies. Shown are immunoblots after detection with anti-HA (^mC^UBC9) and anti-EME1 antibodies. To allow detection of the faint sumoylated forms of ^mC^UBC9 (S* ^mC^UBC9), membranes were cut (indicated by scissors) before separate antibody incubation. **(f)** Model how sumoylation of Ubc9 and EME1 improves rDNA repair. UBC9 and EME1 co-localize in nucleolar repair condensates, which is promoted by their sumoylation. CPT induced seDSBs cannot be resolved by NHEJ as they lack the second end for ligation.

Therefore, seDSB rely on repair by homologous recombination, which is error-prone in repetitive DNA. EME1 sumoylation enhanced by S*UBC9 promotes EME1-MUS81 dependent resolution (red dotted line) of nicked Holliday Junctions (HJ) resembling structures that arise upon template switching and D-loop formation at seDSBs. This prevents detrimental second template switches in repetitive DNA and enables completion of DNA replication by the converging replication fork (adapted from^23^).

To understand whether sumoylation regulates the recruitment of UBC9 to DNA lesions we performed laser microirradiation-induced DNA damage tracks in the established ^mC^UBC9-variant expressing cells lines. The recruitment of tagged proteins to the DNA damage tracks was followed over time. Indeed, both proteins were efficiently recruited to the DNA lesions, but ^mC^S^~^UBC9 showed a faster recruitment and stronger signal at DNA lesions over time than ^mC^UBC9 (Fig. 4c).

To sum up, our data suggests that UBC9 is recruited to sites of DNA damage in the nucleolus in a manner promoted by its sumoylation.

### S^~^UBC9 and EME1 co-localize in nucleolar repair condensates

We next asked whether we could detect co-localization of ^mC^UBC9-variants with EME1. To this end, we ectopically expressed ^GFP^EME1 in the ^mC^UBC9-variant cell lines, exposed the cells to the different treatments, and removed soluble proteins with CSK buffer before immunofluorescence staining. As shown in Fig. 4d and Extended Data Fig. 4c, ^GFP^EME1 was already detected in mock treated cells in fuzzy, ML792-sensitive, speckles and weak colocalization was only detected in the presence of ^mC^S^~^UBC9. Exposure to CPT resulted in enrichment of ^GFP^EME1 in these nucleolar repair condensates, accompanied by a change in shape towards rounding and co-localization with ^mC^UBC9 and ^mC^S^~^UBC9. ML792 pretreatment abolished co-localization with ^mC^UBC9 upon CPT but reduced it only with ^mC^S^~^UBC9, indicating synergistic functions of ^mC^S^~^UBC9 and ^GFP^EME1 in the nucleolus (Fig. 4d).

We then asked whether we would be able to biochemically monitor the shift of UBC9 from the soluble to the insoluble state by cell fractionation and detect changes in UBC9 sumoylation in a CPT- and ML792-regulated manner. Because all UBC9 antibodies tested were not sensitive enough to detect endogenous UBC9 sumoylation, we relied on the ^mC^UBC9 expressing cell line. We used RIPA buffer extracts as the soluble fraction (Supernatant) and dissolved the RIPA insoluble pellets (Pellet) under harsher conditions with 0.5% SDS and sonication before diluting the samples for immunoprecipitations (IP) with anti-HA antibodies. We also compared mock, CPT and ML792 & CPT treated cells. To prevent the large amount of unmodified ^mC^UBC9 from interfering with detection of the sumoylated form, we cut the membranes between the two forms and incubated them individually with anti-HA antibodies. Using this procedure, we were indeed able to detect a slower migrating form of ^mC^UBC9 in mock and CPT treated cells (Fig. 4e). Based on the size shift and sensitivity to ML792 treatment, this band most likely represents the sumoylated form of ^mC^UBC9. Because sample preparation is performed under reducing conditions, we can exclude thioester-bound SUMO for ^mC^UBC9 (2% SDS, 33 mM DTT in Laemmli buffer). Of note, we did not observe an increase in ^mC^UBC9 sumoylation upon CPT treatment. However, the sumoylated ^mC^UBC9 decreased in supernatant and accumulated in the pellet fraction (Fig. 4e), where it interacts with ^GFP^EME1 upon CPT exposure in a ML792 regulated manner, further supporting a cooperative function of ^mC^UBC9 sumoylation and ^GFP^EME1 in nucleolar repair-condensates.

In conclusion, our data suggest that co-localization of UBC9 and EME1 in nucleolar repair-condensates is promoted by CPT treatment and UBC9 sumoylation.

## Discussion

In this study we shed light on the biological function of UBC9 sumoylation upon CPT treatment. Examination of a UBC9 sumoylation mimetic fusion (^mC^S^~^UBC9) revealed that sumoylation promotes the stability and nuclear localization of UBC9 and is beneficial for cell survival. We identified and verified EME1, the regulatory subunit of the structure-specific endonuclease EME1-MUS81 complex, as substrate for the sumoylated UBC9. Both EME1 and MUS81 are among the top hits causing resistance to CPT^7^ and they induce DSBs and replication advantageous for cell survival upon CPT exposure^2^. The cooperative biological function of EME1 and ^mC^S^~^UBC9 is supported by the fact that an EME1 sumoylation mimetic fusion is also beneficial to cell survival upon CPT exposure, and that expression of ^mC^S^~^UBC9 promotes DSB formation and replication. Unexpectedly, we found that expression of ^mC^S^~^UBC9 recruits the replication machinery to nucleoli in a CPT-dependent manner. Moreover, we could confirm its co-localization with EME1 in nucleolar repair-condensates supporting synergistic nucleoli-specific repair functions of ^mC^S^~^UBC9 and the EME1-MUS81 complex.

Nucleoli present the sites of ribosome biogenesis and form around ribosomal DNA (rDNA) placed in the nucleolar organizer region (NOR) of the five acrocentric chromosomes. rDNA is highly repetitive, rich in secondary structures and strongly transcribed by DNA Polymerase I. These combined features define rDNA as one of the most fragile loci in the genome that functions as stress sensor for the cell^24,25^. The optimal number of rDNA units is critical to maintain efficient ribosome biogenesis, cellular longevity, overall genome function and cell survival under stress^26^. Not surprisingly, CPT treatment has been shown to impair ribosome biogenesis^27^. CPT induces nicks in DNA that result in single ended DSBs (seDSBs) when the replication fork “runs off’ on the lagging strand. These particular DSBs cannot be repaired by non-homologous end joining (NHEJ) as they lack the second end for ligation. Therefore, seDSB rely on repair by homologous recombination (HR)^28,29^. However, the repetitive nature of rDNA and its presence on multiple chromosomes make recombination-directed repair in rDNA error-prone that can lead to loss of repeats and viability in the 45S rDNA^30^. Homology-directed repair in nucleoli occurs at all stages of the cell cycle and is accompanied by the movement of DSBs from the nucleolar interior to the periphery^31^.

How can UBC9 and EME1-MUS81 sumoylation improve rDNA repair by HR? We propose a model in which seDSBs in the repetitive rDNA perform resection and template switches in cis or in trans for D-loop switching (Fig. 4f). Due to the seDSBs, the resulting structures resemble nicked Holliday Junctions (nHJs), the natural substrates of the EME1-MUS81 complex^32–34^. Resolution of nHJ at this stage is highly beneficial as it limits resection and prevents a second template switch that carries a high-risk of instability particular in repetitive regions. The converging replication fork can then complete DNA replication, comparable to a mechanism described for MUS81 dependent repair of repetitive Alu elements upon replication fork breakage^23^. Further support for our model comes from studies in *Saccharomyces cerevisiae* showing that MUS81 activity is required for the maintenance of rDNA repeat number by functioning in the quality control of replication forks^35^. Due to the fragile nature of ribosomal chromatin, rDNA damage repair is constitutively required, which explains the survival advantage of ^mC^S^~^UBC9-expressing cells already in mock-treated cells.

In addition to new insights into rDNA damage repair, our results reveal a molecular mechanism leading to CPT-based drug resistance. Various studies have shown that UBC9 is upregulated in multiple cancers including brain, ovarian, prostate, lung, liver, skin cancer and B-cell lymphoma^13,14,36^ and high UBC9 levels correlate with poor response to chemotherapy and clinical prognosis^16,37^. Inhibition of sumoylation pathway by targeting the E1 enzyme delays tumor progression^9,10,12,17^. Whether an increase in UBC9 sumoylation also corresponds to an increase in UBC9 levels remains to be shown, but due to the generally very low levels, this is difficult to demonstrate. However, we could envision that high UBC9 levels also induce UBC9 sumoylation because, at least in *in vitro* sumoylation assays UBC9 is sumoylated in the absence of E3 ligases. Remarkably, CPT treatment does not appear to induce UBC9 sumoylation but rather causes its recruitment to nucleolar condensates where it interacts with its substrate EME1. In summary, we could envision that acquired or de novo clinical CPT-based drug resistance is regulated by patient and tumor specific UBC9 expression levels and consequently UBC9 and EME1 sumoylation. We propose that a fine-tuned combinatorial treatment between inhibition of the SUMO pathway and CPT-derived drugs may enable the definition of personalized CPT-based cancer treatments.

## Methods

### Antibodies

Rabbit polyclonal-antibodies were raised by Coring System Diagnostix GmbH against our purified recombinant human GST- and untagged EME1 full-length protein, validated (Fig. 2) and used as serum. anti-GST was a kind gift from Dr. Egon Ogris. All other primary antibodies were purchased and validated by the manufacturers (data available on manufacturers’ websites).

#### Antibodies used in Immunoblot analysis (IB)

mouse anti-UBC9 (SAB 1305501, Sigma Aldrich), mouse anti-GFP (1814460, Roche), mouse anti-SUMO1 (18-2306, Zymed laboratories Inc.), mouse anti-SUMO2/3 (ab81371, Abcam), mouse anti-Lamin B1 (A5316, Sigma-Aldrich), mouse anti-β-actin (A2228, Sigma-Aldrich), mouse anti-HA (Clone 16B12, Covance), rabbit anti-HA (26183, Thermo Fischer). Secondary-antibodies against rabbit (711-035-152, Jackson) or mouse (715-035-150, Jackson) were horseradish-peroxidase conjugated.

#### Antibodies used in IF

anti-HA (26183, Thermo Fischer), mouse anti-mCherry coupled to Alexa 594 dye (M11240, Thermo Fischer), goat anti-GFP FITC (NB100-1771, Novus Biologicals), mouse anti-γH2AX antibody (05-636, Millipore). Secondary antibodies for detecting HA was wanti-mouse coupled to Alexa 488 dye (NC1214603, Jackson) and for γH2AX was anti-mouse coupled to Alexa 594 dye (Jackson, 715-585-150). Nucleolar staining was with Syto RNA select green fluorescent stain (S32703, Thermo Fischer).

#### Antibodies for FACS analysis

anti-γH2AX coupled Alexa 647 (9720, Cell signaling), anti-mCherry coupled to Alexa 594 dye (M11240, Thermo Fischer).

### Plasmids

UBC9 and S^~^UBC9 cDNA were tagged with HA-Strep-mCherry and cloned into the mammalian expression vector pcDNA5 in order to generate stable cell lines in U2OS cells using Flp-In system (^mC^UBC9, as wild-type and the SUMO2 fusion ^mC^S^~^UBC9). Expression constructs for MUS81 and EME1 were generated by PCR amplification from human cDNA. For mammalian replacement studies, EME1 was subcloned into a doxycycline (Dox) inducible pTRE3G vector and siRNA resistance was generated by introducing silent mutations by site directed mutagenesis. MUS81 was subcloned into pCRUZ-Myc. EME1 mutants were obtained, using site directed mutagenesis. HA-Strep-GFP-EME1 (^GFP^EME1) was subcloned in pcDNA5. For bacterial expression, MUS81 and EME1 were cloned into pET23a+ and pET28a+ plasmids, respectively. EME1 K27/28R, ΔSIM1 (54-IVVV-57 to AAAA) and ΔSIM2 (79-VRLL-82 to AAAA) were generated by site directed mutagenesis. For antibody raising, EME1 was cloned into pGEX6P1. The S2^~^UBC9 linear fusion was cloned into a pGEX6P1 expression vector by restriction free cloning^1^. All constructs generated were verified by sequencing at the MPI-IE DNA Sequencing Core Facility.

### siRNA

siRNA for the replacement studies was ordered from Amsbio (EME1: #SR315576A, scrambled siRNA: #SR30004).

### Cell lines & culture

U2OS (Human bone osteosarcoma epithelial cells) were grown in McCoy’s 5A (Modified) medium (16600082, Thermo Fischer) supplemented with 10% fetal bovine serum (FBS) (F0804, Sigma-Aldrich) and Penicillin-Streptomycin (P4458, Sigma-Aldrich). Hygromycin (CP12.2, Carl Roth), blasticidin (15205, Sigma-Aldrich) and Tet System approved FBS (631106, Takara) containing medium was used to culture Dox (A2951, PanReac Applichem) induced stable cell lines. Adherent HeLa cells, a kind gift from Lea Sistonen, were grown in (Dulbecco’s Modified Eagle’s Medium (DMEM) (D5796, Sigma-Aldrich), 10 % 26140079, fetal bovine serum (FBS) (Thermo Fisher), 1 x GlutaMAX (35050061, Thermo Fisher), 1 x Penicillin-Streptomycin (15140122, Thermo Fisher). The adherent HeLa cells were mycoplasma tested before stock freezing. Subconfluent cultures were maintained at 37°C in a humidified incubator with 5% CO_2_ and 95% air atmosphere. Cells were used without further authentication.

U2OS cells modified with the Flp-In TRex system were a kind gift from Dr Jakob Nilsson. ^mC^UBC9 and ^mC^S^~^UBC9 stable cell lines were generated in U2OS cells according to manufacturer’s instructions (Flp-In TRex manual, Invitrogen). Single clones were verified for inducible expression of the appropriate ^mC^UBC9 variant, via PCR, western blotting, and microscopy.

U2OS cells expressing His-SUMO2 were described in^2^. U2OS cells expressing His-SUMO1 were construct by lentiviral infection at MOI=2 of a construct expressing 10xHis-N-terminal tagged-SUMO1 where all lysines had been mutated into arginines. After infection, cells were selected with puromycin (A1113802, Thermo Fisher) at 3 μg/mL.

### Immunofluorescence assay (IFA)

^mC^UBC9 and ^mC^S^~^UBC9 cell lines were seeded in Nunc Lab-Tek chambered slides and cells were either transfected (for 14 hours) and/or induced by 30 and 10 ng/ml Dox, respectively (for 16 hours) in Tet FBS medium. Subsequent cells were treated by DMSO (vehicle control), Camptothecin (CPT) (10 μM for 1 hour) (C9911, Sigma-Aldrich), ML (ML792) (1 μM for 2 hours) (gift from UbiQ) and ML & CPT (ML was added an hour before addition of CPT). After treatments, cells were washed with 1X PBS (14190250, Thermo Fischer), followed by fixation with 4% formaldehyde (F8775, Sigma-Aldrich) for 15 minutes and permeabilization with 0.5% triton X 100 (3051, Carl Roth) for 10 minutes. Coverslips were then blocked with Sea-block buffer (37527, Themo Fischer) for 1 hour at room temperature, washed with 0.2% triton X 100 in 1x PBS and incubated with primary antibodies (1:300 dilution in Sea-block buffer) overnight at 4°C. Further washing with 0.2% triton X 100 in 1x PBS, was followed by a one-hour incubation with appropriate secondary antibodies at room temperature (1:1000 dilution in Seablock buffer). The slides were washed again before being fixed with anti-fade DAPI (S36939, Thermo Fischer). Primary antibodies used were either anti-HA (26183, Thermo Fischer), anti-mCherry coupled to Alexa 594 dye (M11240, Thermo Fischer) and anti-GFP FITC (NB100-1771, Novus Biologicals). Secondary antibody for detecting HA was anti-mouse coupled to Alexa 488 dye (NC1214603, Jackson). Images were acquired with a Zeiss LSM780 confocal microscope using 63x oil immersion objective at a zoom range 0.6-2x. Images were processed using Zen black & blue or Imaris software.

For EdU and γH2AX staining, CPT and ML treated cells were incubated with 100 μM EdU for 1 hour. Cells were washed, fixed and permeabilized followed by incubation with Click iT Plus Edu reagent as per manufacturer’s protocol (Click-iT™ Plus EdU Kit for Imaging, Alexa Fluor™ 488 dye, C10637, Invitrogen). Subsequently cells were stained as described above with anti-γH2AX antibody (05-636, Millipore) overnight at 4°C and subsequent with anti-mouse coupled to Alexa 594 dye (715-585-150, Jackson,). Slides were fixed using anti-fade DAPI.

### Transfection for co-localization studies (Fig. 4D)

HA-Strep-GFP-EME1 (^GFP^EME1) transfections were carried out in ^mC^UBC9 and ^mC^S^~^UBC9 cell lines using Lipofectamine 3000 transfection reagent (3000-008, Invitrogen) as per manufacturer’s instructions for 24 hours than proceeded with Immunofluorescence assay.

### Analysis of nuclear/nucleolar condensates by pre-extraction with CSK-buffer

Pre-extraction to release soluble proteins and enrich in nuclear condensates was performed by preincubation with CSK buffer as described in ^3,4^ In brief, after seeding and treatments of cells as described above, coverslips were washed and incubated with CSK buffer (10 mM PIPES with pH 7.0, 100 mM NaCl, 300 mM sucrose, 3 mM MgCl2 and 0.7% Triton X-100) for 3 minutes at room temperature. Subsequently, cells were washed, fixed with 4% paraformaldehyde (PFA) (SC281692, Santa Cruz) in PBS for 25 minutes at room temperature before staining the cells with the respective antibodies as described above.

For nucleolar staining, cells were treated with CSK buffer and fixed (as per CSK buffer protocol). Coverslips were washed and incubated for 20 minutes with 5μM Syto RNA select green fluorescent stain (S32703, Thermo Fischer) and washed thrice with 1X PBS. Slides were fixed using anti-fade DAPI.

### Flow cytometry

^mC^UBC9 and ^mC^S^~^UBC9 cell lines were seeded in 10 cm dishes and induced with 30 and 10 ng/ml Dox respectively. Cells were treated with 1 μM ML792 for 2 hours and 10 μM CPT for 1 hour, followed by EdU pulse (10 μM) for 30 minutes. Trypsinized cells were washed with 1% BSA in PBS, and fixed with 4% paraformaldehyde (PFA) for 15 minutes at room temperature. After additional washing, cells were permeabilized with the 1X Saponin based solution provided in Click iT Kit (C10632, Thermo Fischer) for 15 minutes and treated with Click-iT Plus EdU reaction mix as per manufacturer’s instructions (Click-iT Plus EdU Alexa Fluor 488 Flow Cytometry, C10632, Thermo Fischer). Samples were incubated at room temperature for 30 minutes, washed with 1X Saponin based wash buffer, blocked with Sea-block buffer (37527, Thermo Fischer) for one hour at room temperature and co-stained with anti-γH2AX coupled to Alexa 647 (560447, BD Biosciences) and anit-mCherry coupled to Alexa 594 (M11240) antibodies for overnight at 4°C in the dark. DAPI (D1306, Thermo Fischer) was added to a final concentration of 1 mg/ml to facilitate exclusion of dead cells from the analysis for 15 minutes at room temperature and cell pellets were resuspended in 500 μl of cold PBS. Samples were analyzed with a Fortessa III flow cytometer (BD Biosciences) and FlowJo software. Graphs were plotted using Graphpad Prism 8.0.

### Laser microirradiation

Laser track experiments were performed using a Leica SP5 confocal microscope with an environmental chamber set to 37 °C. Briefly, ^mC^UBC9 and ^mC^S^~^UBC9 cell lines were grown on 18 mm coverslips and media was replaced by fresh medium with 20 μg/mL Dox. Prior to micro irradiation, medium was replaced by CO_2_-independent Leibovitz’s L15 medium supplemented with 10% FCS, Penicillin-Streptomycin and 20 μg/mL Dox. Laser micro-irradiation was carried out on a Leica SP5 confocal microscope equipped with an environmental chamber set to 37°C. DNA damage tracks (1 μm width) were generated with a Mira modelocked titanium-sapphire (Ti:Sapphire) laser (l = 800 nm, pulse length = 200 fs, repetition rate = 76 MHz, output power = 80 mW) using a UV-transmitting 63 × 1.4 NA oil immersion objective (HCX PL APO; Leica). Gain was set at 30 % and offset at 50 %, output was 1.5 ± 0.5 W. Relative recruitment values were analyzed using Leica LAS X Software.

### Clonogenic survival assays

For transfection, U2OS cells were seeded and EME1 knockdown was performed using siRNA of EME1 with Lipofectamine RNAiMAX transfection reagent (LMRNA015, Invitrogen) according to manufacturer’s instructions. Cells were transfected with indicated plasmids (empty vector, wild type EME1 (WT), EME1-K27R (KR) mutant, and S1^~^EME1 in pTRE3G vector) using Lipofectamine 3000 transfection reagent. Dox concentrations for induction were adjusted to obtain expression levels comparable to the endogenous EME1 protein (~2 ng/ml Dox). After 48 hours post-transfection, cells were treated with CPT (0, 10, 100, 500 nM) for 2 hours, DMSO was used as a control in all the experiments.

Similarly, ^mC^UBC9 and ^mC^S2^~^UBC9 cell lines were seeded in 6 well plate and induced with Dox (30 ng/ml and 10 ng/ml respectively) for 16 hours. Cells were treated with CPT (0, 10, 100 and 500 nM) for 1 hour. Subsequently, cells were washed with 1X PBS, trypsinized and seeded in triplicates into 6-well plates at a density of 500 cells per well. After 7-12 days post CPT treatment, the cells were washed, fixed with PFA and stained with 0.5% Crystal Violet solution (V5265, Sigma Aldrich) in 20% methanol and photographed on a light table. The images were analyzed with ImageJ and colonies were counted for each well. Plots and statistics were performed by two-way ANOVA analysis using Graphpad Prism.

### Cell extracts for Immunoblotting (IB)

Dox induced cells were harvested and lysed in lysis buffer to proceed for Western blotting (50 mM Tris-Cl pH-6.8, 0.1% bromophenol blue, 10% glycerol, 100 mM DTT, 5 mM EDTA, 2% SDS). Lysed protein samples were sonicated and resolved by SDS-PAGE. Proteins were transferred onto the nitrocellulose membrane using semi-dry transfer apparatus. Eventually, the blots were probed with indicated primary antibodies: anti-HA (261883, Thermo Fischer) and anti-β-actin (A2228, Sigma-Aldrich) antibodies overnight at 4°C followed by incubation with secondary anti-rabbit (711-035-152, Jackson) or anti-mouse (715-035-150, Jackson) antibodies for 1 hour at room temperature. Detection was with ECL reagent (1705061, Biorad).

### Immunoprecipitation

^mC^UBC9 and ^mC^S^~^UBC9 cell lines were grown in 1 × 15 cm dishes and induced with Dox (1 μg/ml). Cells were treated with 1 μM ML for 2 hours, 10 μM CPT for 1 hour and PR619 for 30 minutes (PR619 blocks ubiquitin and SUMO proteases, 16276, Cayman Chemical). Cells were scraped and lysed in RIPA buffer (20mM Tris pH 7.5, 150mM NaCl,5mM EDTA, 1% NP40, 5% glycerol, 0.5% Na-deoxycholate, 0.1% SDS) supplemented with protease inhibitors (1 mM Leupeptin(A2183, Merck), Pepstatin A (A2205, Merck), 1 mM Aprotinin (A1623, Roth), 100 mM Pefabloc (11873601001, Merck), 400 mM Iodoacetamide (I1149, Sigma), 400 mM N-Ethylmaleimide (E1271, Sigma) and 200 mM Orthovanadate (P07581, New England Biolabs)). Lysed cells were sonicated, and then separated in supernatant (RIPA buffer soluble fraction) and pellet fractions. Pellets were dissolved in RIPA buffer containing 0.5 % SDS, then lysed, by additional sonication before further dilution with buffer (20 mM Tris pH 7.5 and 150 mM NaCl along with protease inhibitors as mentioned above) to reduce SDS concentration from 0.5 % to 0.1 % as it is required for IPs (RIPA buffer pellet fraction).

Subsequent immunoprecipitations were performed from both fractions by incubating lysates with anti-HA antibodies (mouse monoclonal, Clone 16B12, Covance) overnight at 4°C. Immunoprecipitates were collected on pre-equilibrated (RIPA buffer) protein A Sepharose beads (53139, Thermo Scientific) by incubating the reaction for 2 hours at room temperature on a rotating wheel. After additional washing steps with RIPA buffer, beads were boiled in 2X Laemmli buffer for 5 minutes at 95°C. Samples were resolved by 7% SDS-PAGE and transferred to nitrocellulose membrane by semi-dry i-blot transfer system (IB1001, Invitrogen). Inputs and elution fractions were analyzed on separate gels. For detecting the sumoylated form of UBC9, blots was cut between 70 and 55 kDa and probed separately. Blots were incubated with primary rabbit polyclonal anti-HA (26183, Thermo Fischer) or anti-EME1 (homemade) antibodies at 1:2000 dilutions overnight at 4°C, followed by incubation with anti-rabbit secondary antibodies (711-035-152, Jackson) at 1:10.000 dilution for 1 hours at room temperature. Detection was with ECL reagent (1705061, Bio-Rad) on a Bio-Rad ChemiDoc imaging system (17001402, Bio-Rad).

### His-SUMO pull downs

His-tagged SUMO conjugates were purified from U2OS cells as described before ^2^. Briefly, U2OS cells stably expressing His-SUMO were washed and collected in PBS. Input samples were taken and the cells were lysed in Guanidinium lysis buffer. After sonication, the lysates were equalized using BCA Protein Assay Reagent (Thermo Scientific). Subsequently, the His-SUMO conjugates were enriched using nickel-nitrilotriacetic acid beads (Qiagen). Upon multiple washing steps, the His-SUMO conjugates were eluted from the beads and separated by SDS-PAGE.

### Protein purification

All bacterial expression constructs were transformed into chemically competent bacteria, cultured in lysogeny broth (LB) and selected with the appropriate antibiotics.

Purification of mammalian SUMO1, SUMO2, SUMO E1 (Aos1/Uba2), SUMO E2 (Ubc9), GST-RanBP2ΔFG, 6xHis-PIAS1, 6xHis-MPB-ZNF451-N, 6xHis-MPB-ZNF451-3, 6xHis-MPB-KIAA1586, has been described ^5–8^. S1*UBC9 and S2*UBC9 was purified as described ^6^. S2^~^UBC9 expression was performed in BL21 gold strains in LB. Bacterial culture waw grown at 37 °C to a density (OD600) ≈ 0.5. Protein expression was induced with a final concentration of 1 mM IPTG for 4 hours. Subsequently, cells were harvested in 50 mM Tris, pH 8.0, 200 mM NaCl, 1 mM dithiothreitol (DTT) buffer supplemented with protease inhibitors (1 μg/ml of leupeptin, 1μg/ml pepstatin, 1 μg/ml aprotinin and 1 mM phenylmethylsulfonyl fluoride) and lysed by freeze-thawing and incubation with 1 mg/ml lysozyme for 30 minutes on ice. Lysates were clarified by centrifugation (31,200 RCF for 30 minutes), and the supernatant was subjected to batch affinity purification with Glutathione Sepharose 4B Resin (GE Life Sciences). Bound protein was eluted in harvesting buffer containing GST-Precission protease, by incubating over night at 4° C. The purified S2^~^UBC9 was concentrated and subjected to size-exclusion chromatography (SEC) with a Superdex 75 10/300 GL column (GE Life Sciences) on an ÄKTA explorer fast protein liquid chromatography (FPLC) system (GE Life Sciences) in 20 mM HEPES, 110 mM K-acetate, 2 mM Mg-acetate, 0.5 mM EGTA, 1 mM DTT supplemented with protease inhibitors (1 μg/ml of leupeptin, 1μg/ml pepstatin and 1 μg/ml aprotinin). Fractions that contained the desired protein were pooled and flash frozen.

MUS81-6x-His-EME1 expression was performed in Rosetta(DE3) strains in LB. Bacterial culture was grown at 37 °C to a density (OD600) ≈ 0.5. Protein expression was induced with a final concentration of 0.1 mM IPTG over night at 16° C. Subsequently, cells were harvested in 50 mM sodium-phosphate, pH 6.5, 100 mM NaCl, 100 mM imidazole, 10 % glycerol, 0.1 % Triton X 100, 1 mM β-mercaptoethanol (β-ME) buffer supplemented with protease inhibitors (1 μg/ml of leupeptin, 1μg/ml pepstatin, 1 μg/ml aprotinin and 1 mM phenylmethylsulfonyl fluoride) and lysed by freeze-thawing and incubation with 1 mg/ml lysozyme for 30 minutes on ice. Lysates were clarified by centrifugation (31,200 RCF for 30 minutes), and the supernatant was subjected to batch affinity purification with ProBond Nickel-Chelating Resin (Life Technologies). Bound protein was eluted in harvesting buffer containing 500 mM imidazole. The purified MUS81-EME1 was subjected to cation-exchange chromatography (CEC) with a Mono S 4.6/100 PE column (GE Life Sciences) on ÄKTA explorer FPLC system. The protein was eluted in a 3 step gradient between buffer A (50 mM sodium-phosphate, pH 6.5, 100 mM NaCl, 10 % glycerol, 0.1 % Triton X 100, 1 mM DTT supplemented with protease inhibitors (1 μg/ml of leupeptin, 1μg/ml pepstatin, 1 μg/ml aprotinin) and buffer B (buffer A with 1000 mM NaCl). Step 1 was for 5 column volumes (CV) at 20 % buffer B, Step 2 was for 5 CV at 60 % buffer B and step 3 was for 5 CV at 100 % buffer B. Fractions that contained the desired protein were pooled and concentrated. Subsequently the buffer was exchanged to 20 mM HEPES, 110 mM K-acetate, 2 mM Mg-acetate, 0.5 mM EGTA, 100 mM NaCl, 10 % glycerol, 0.1 % Triton X 100, 1 mM DTT supplemented with protease inhibitors (1 μg/ml of leupeptin, 1μg/ml pepstatin and 1 μg/ml aprotinin) using a PD MidiTrap G-25 buffer exchange column and the protein was flash frozen.

### *In vitro* sumoylation assay

The assays were performed as described before^5–8^. Briefly, reactions were performed in 20 μl volumes containing 20 mM HEPES pH 7.3, 110 mM potassium acetate, 2 mM magnesium acetate, 0.05% (v/v) Tween20, 1 mM DTT, 0.2 mg/ml ovalbumin, and 5 mM adenosine triphosphate (ATP). Samples were incubated at 30°C for 30 minutes. Reactions were terminated by boiling with 2X Laemmli buffer at 95°C for 5 minutes and samples were resolved by SDS-PAGE. Detection was done as indicated.

### Protein array screen

*In vitro* sumoylation reactions were performed on protein array slides with more than 9000 human proteins spotted (ProtoArray® Human Protein Microarray v5.0, Invitrogen). The reaction was performed by the technical service of Invitrogen with our purified enzymes, buffers following our protocol (see above). Instead of duplicates, two E2 enzyme concentrations were used. In brief, the array slides were incubated for 1hour at 30°C with a reaction mix containing 4 μM HA-SUMO1, 100 nM E1, an ATP regenerating system and either UBC9 or S1*UBC9 each at two different concentrations (100 nM or 500 nM). Two arrays were included as controls, one with a reaction mix without E2 enzyme and one with only the detection reagents. Detection was with anti-HA antibody and subsequently with Alexa 647-labelled secondary antibodies. After final washing steps, the arrays were dried and scanned.

### Statistics

GraphPad Prism v8.0 (GraphPad Software) was used to plot and analyze data. All experiments with quantitative analysis included data from at least three independent experiments. Data were expressed as mean ± SEM, and statistical differences were calculated by two-way ANOVA using Tukey test for grouped samples.

## Reporting summary

Further information on research design is available in the Nature Research Reporting Summary.

## Data availability

No source data are provided with this manuscript.

## Acknowledgements

Our special thanks go to all former and current members of A.P.’s laboratory for discussions and sharing reagents and the MPI-IE core facilities for Proteomics, Imaging, and Flow cytometry for their technical help. ML792 was a kind gift from Farid El Oualid and Alfred Nijkerk (UbiQ, Amsterdam). For sharing reagents, we kindly acknowledge Lea Sistonen, Jakob Nilsson and Egon Ogris. The Pichler lab is currently funded by the Max Planck Society and the European Union’s Horizon 2020 research and innovation program under the Marie Sklodowska-Curie grant agreement 765445. Parts of this work were funding by the WWTF Project Number LS05003 and the DFG PI 917/1-1 in the past. Román González Prieto was supported by the Dutch Cancer Society, (KWF-YIG 11367 / 2017-2). This article is based on work from European Cooperation in Science and Technology (COST) Action (PROTEOSTASIS BM1307 and ProteoCure CA20113 to A.P and A.C.O.V.) and, supported by COST.

## Author contribution

J.B and A.P. initiated the project and A.P. has set the direction of this study. J.B., E.N., K.B., R.G.P and A.P. designed the majority of experiments. C.F. generated the stable inducible UBC9 cell lines. J.B. cloned and expressed the recombinant EME1 and MUS81 and performed EME1 replacement studies. In vitro sumoylation assays were performed by J.B. N.E and E.N.; V.K.C. purified multiple proteins for the in vitro sumoylation assays; clonogenic survival assays by J.B and E.N; FACS analysis by E.N; all IFAs and expression analysis of UBC9 variant cell lines by K.B; recruitment assays and the in vivo EME1 sumoylation analysis by R.G.P.; statistical analysis by E.N., K.B., R.G.P.; in vitro sumoylation on Proteinarray by M.G.; UBC9 IPs by E.N. A.P. supervised. K.B., E.N., J.B., C.F., N.E. and A.C.O.V. supervised R.G.P.; K.B., E.N., J.B., C.F., N.E., and R.G.P. prepared Fig.s, Fig. legends and wrote the methods part, A.P. wrote the manuscript with the input from all authors.

---

**Extended Data Fig. 1.**
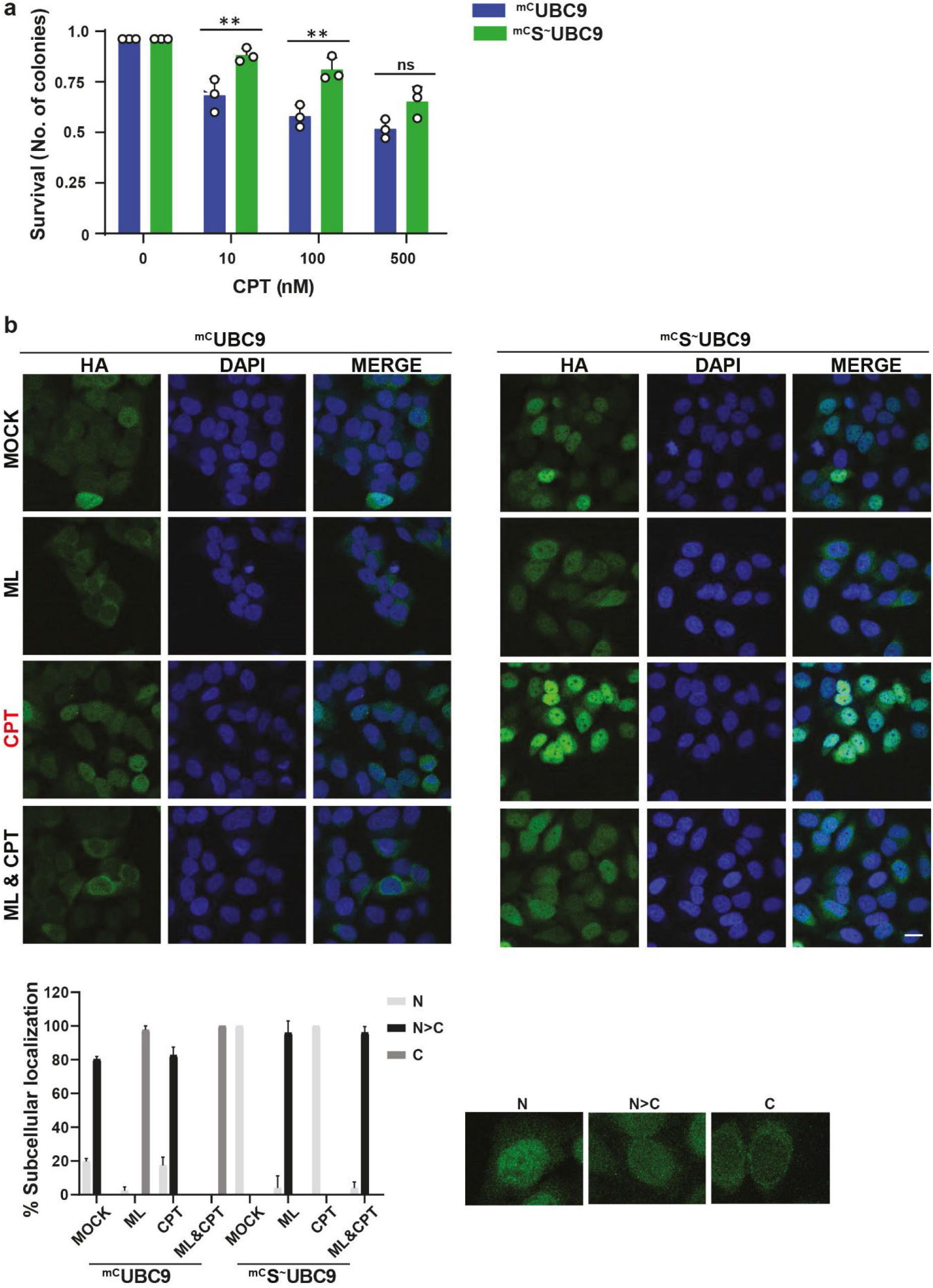
Sumoylation promotes UBC9’s stability, nuclear localization and cell survival. Related to Fig. 1. **(a)** Diagram of clonogenic survival assay related to Fig. 1a but normalized to mock conditions. **(b)** Multi-cell images related to Fig. 1e. Graph shows the average values and error bars (+SEM) of ^mC^UBC9 and ^mC^S^~^UBC9 subcellular localization upon different treatments in percentage from three biological experiments (N=50-100 cells/experiment). N and C represent Nuclear and Cytosolic localization, respectively.

**Extended Data Fig. 2.**
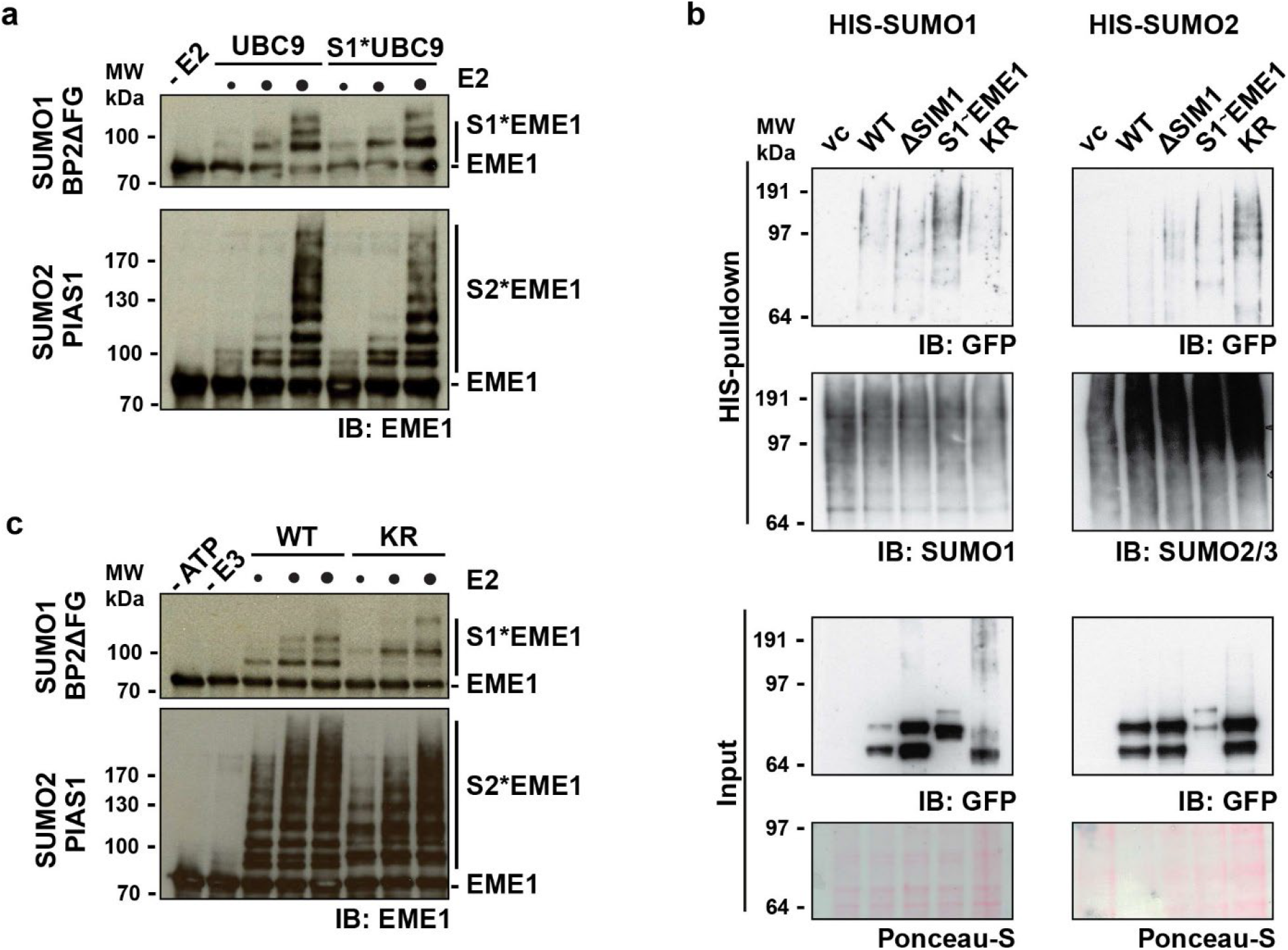
EME1 is a S*UBC9 substrate and its sumoylation also promotes cell survival. Related to Fig. 2. **(a)** Immunoblot of *in vitro* sumoylation assays as in Fig. 2b, but E3-independent reaction with increasing concentrations of Ubc9 or S2-Ubc9 (62.5, 125, 250 and 500 nM) in the presence of 50 nM Aos1-Uba2, 200nM EME1 WT and 2 μM SUMO1 (upper panel) or 2 μM SUMO2 (lower panel). **(b)** Immunoblots of His-SUMO purifications from U2OS cells expressing either His-SUMO1 (left panel) or His-SUMO2 (right panel). Cells were transiently transfected with indicated GFP-EME1 variant or vector control (vc). 2 days after transfection, His-SUMO conjugates were purified and analyzed. **(c)** Immunoblots as in Fig. 2b but with increasing concentrations of Ubc9 or S1*Ubc9 (1, 5 and 25 nM) in the presence of 50 nM Aos1-Uba2, 200nM EME1 WT, 2 μM SUMO1 and 5 nM BP2ΔFG (upper panel) or 2 μM SUMO2 and 100 nm PIAS1 (lower panel).

**Extended Data Fig. 3.**
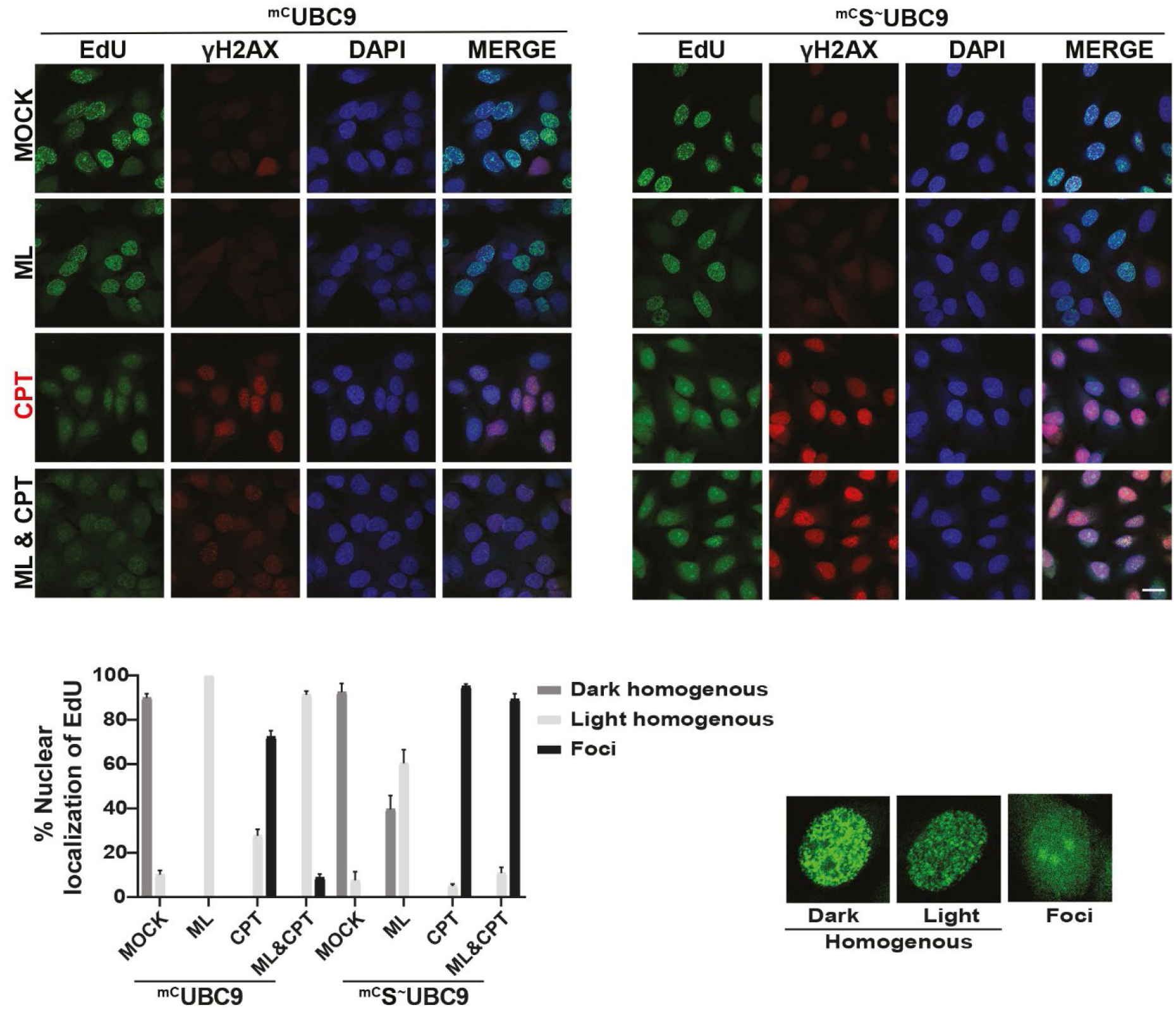
S^~^UBC9 expression induces DSBs and replication upon CPT exposure. Related to Fig. 3. Multi-cell images related to Fig. 3c. Graph shows the average values and error bars (+SEM) of the different nuclear localizations of EdU in ^mC^UBC9 and ^mC^S^~^UBC9 expressing cells upon different treatments in percentage of three biological replicates (N=50-100 cells/experiment).

**Extended Data Fig. 4.**
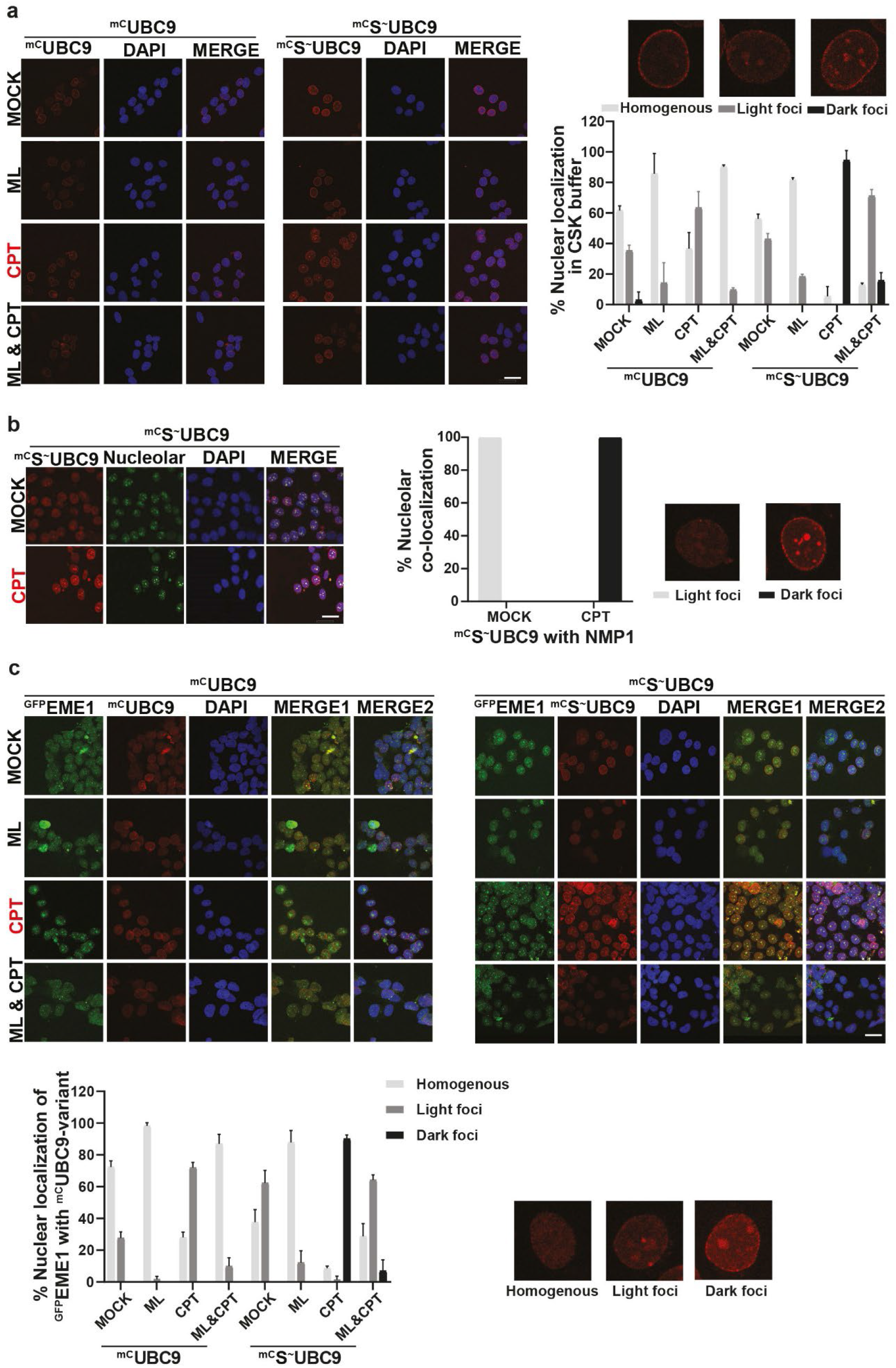
S^~^UBC9 and EME1 co-localize in the nucleolus. Related to Fig. 4. **(a)** Multi-cell images related to Fig. 4a. Graph shows the average values and error bars (+SEM) of the different nuclear localization of ^mC^UBC9 and ^mC^S^~^UBC9 upon indicated drug exposure and CSK buffer treatment in percentage of three biological replicates (N=50-100 cells/experiment). **(b)** Multi-cell images related to Fig. 4b. Graph shows the average values and error bars (+SEM) of the nucleolar localization of ^mC^S^~^UBC9 upon different treatments in percentage of three biological replicates (N=50-100 cells/experiment). **(c)** Multi-cell images related to Fig. 4d. Graph shows the average values and error bars (+SEM) of the nucleolar localization of ^GFP^EME1 with ^mC^UBC9-variants upon different treatments in percentage of three biological replicates (N=50-100 cells/experiment).

**Extended Data Fig. 5.**
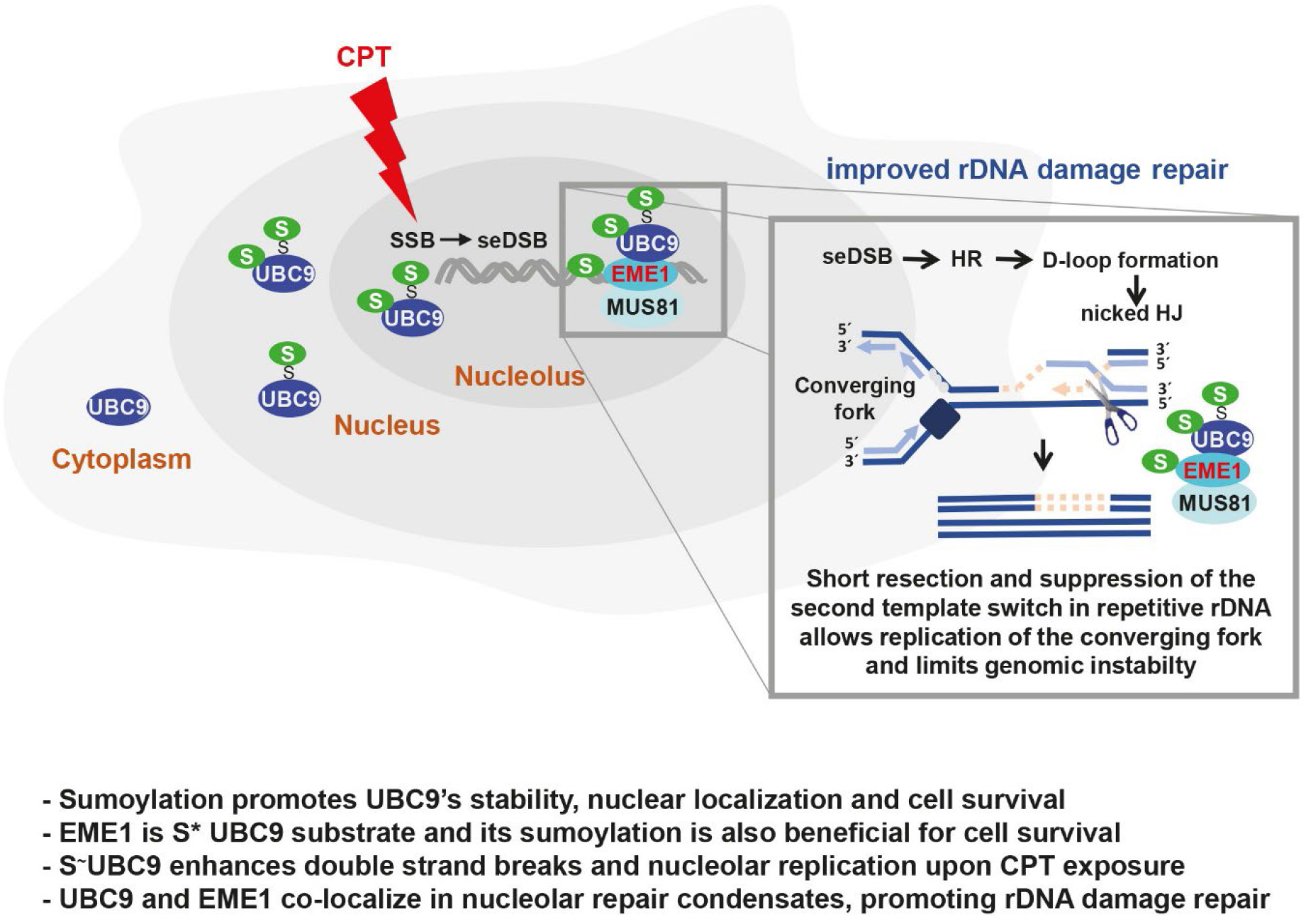
Model. Key findings of this study are summarized in bullet points. More detailed information in the text.

